# The first complete assembly for a lungless urodelan with a “miniaturized” genome, the Northern Dusky Salamander (Plethodontidae: *Desmognathus fuscus*)

**DOI:** 10.1101/2024.04.30.591895

**Authors:** Edward A. Myers, R. Alexander Pyron

## Abstract

Salamanders have some of the largest genomes among animals. However, there is a great disparity in total size, ranging from ∼8–120GB depending on the lineage. Species in the lungless genus *Desmognathus* (Plethodontidae) are among the smallest, with estimated genome sizes of 13–15GB. Salamander genomes have exceptional interest in numerous topics ranging from genome-size evolution, the genetic basis of evolutionary differences in life history, and the physiological basis of regeneration, vision, and immunity. However, their large genomes have limited previous attempts at sequencing and assembly, particularly given the difficulties of mapping extensive repeat regions with short-read data. Here we assemble a draft genome of *Desmognathus fuscus* using PacBio HiFi reads and generate transcriptomic data from two specimens. The combined assembly resulted in a 16.1GB genome in 19,640 contigs and an N/L50 of 2,455/1.75MB, with the longest contig at 27.9MB. The assembly and transcriptome are nearly complete with 93% of the 5,310 BUSCO tetrapod orthologs identified. Attempts to scaffold these data to the existing *Ambystoma* genome resulted in only 5.8GB of the *D. fuscus* genome mapping to this reference. This low success suggests substantial syntenic and sequence divergence across salamanders, which may be the result of significant miniaturization in the *Desmognathus* genome. Identification and annotation of transposable elements reveals that only 26% of the genome is single copy with 74% corresponding to TEs. The most common class of TE in the genome are LTRs (35% of the total genome) and LINEs (∼15% of the genome). The relative divergence landscape of these TEs shows an early expansion and slow contraction of LINEs, followed by a quick recent expansion of both LTRs and DNA transposons. This assembly will serve as an important reference for amphibian genomics.

## Introduction

Salamanders have some of the largest genomes known among animals, ranging from ∼8–120GB (Sclavi and Herrick 2019). Among these, significant decreases in genome size are observed in the lungless family Plethodontidae (Sessions and Larson 1987), with a major downshift in the eastern Nearctic genus *Desmognathus* (∼13–15GB; Liedtke *et al*. 2018).

Genome-size changes in salamanders are associated with ecological constraints and life-history strategies (Lertzman-Lepofsky *et al*. 2019), and *Desmognathus* notably represent a reversion to bi-phasic lifestyle with metamorphosing aquatic larvae (Chippindale *et al*. 2004). Consequently, the *Desmognathus* genome is of exceptional interest both for the broad applicability of salamanders as a model system for traits such as regeneration (Sessions and Wake 2021), but also for the evolutionary linkage between repeat elements, ecological adaptations, and genome size (Jockusch 1997; Sun *et al*. 2012b; Itgen *et al*. 2022; Mueller *et al*. 2023).

Previous studies using a short-read shotgun-sequencing approach to generate partial genomic coverage in *Desmognathus* and relatives showed that up to 30% of the genome consists of long terminal repeat (LTR) transposons, primarily Ty3/*gypsy* elements at up to 20% of the reads for each species (Sun *et al*. 2012b). Additionally, these species show extremely slow rates of DNA loss (<0.05 bp/substitution) in these elements, suggesting a primary mechanism for genome expansion in salamanders (Sun *et al*. 2012a). Those authors also noted that their results for LTR content were likely to be substantial underestimates, emphasizing the need for more complete and contiguous assemblies to characterize the repeat-element landscape in plethodontids and other salamanders (Sun and Mueller 2014).

Consequently, salamander genomes are of extensive interest for numerous topics including the genetic basis of evolutionary differences in life history and genome size (Lertzman-Lepofsky *et al*. 2019; Mueller *et al*. 2023), as well as regeneration, vision, and immunity (Roth *et al*. 2008; Joven *et al*. 2019; Bolaños-Castro *et al*. 2021). Yet, their gigantic genomes have limited attempts at assembly (Pyron *et al*. 2024), given the difficulties of mapping extensive repeat landscapes with short-read data. The first species sequenced was the laboratory model axolotl (Ambystomatidae: *Ambystoma mexicanum;* Keinath *et al*. 2015; Evans *et al*. 2018b, 2018a), yielding exceptional insight into several pertinent questions in gene regulation, transcriptional control, and regeneration (Schloissnig *et al*. 2021). In particular, up to 19GB (∼60%) of the 32GB genome may be repeat elements (Keinath *et al*. 2015).

Another genome has recently been made available for a newt (Salamandridae: *Pleurodeles waltl*), comprising a 20GB chromosome-scale assembly providing similarly extensive insight into gene complexes related to regeneration and showing exceptional diversity of repeat elements (Elewa *et al*. 2017; Brown *et al*. 2022). Similarly, recent authors produced an unscaffolded assembly for a second newt (*Calotriton arnoldi*) based on pooled reads from two captive-bred siblings, with ∼82% completeness for the BUSCO Vertebrata dataset (Talavera *et al*. 2024). While single-individual genomes remain the gold standard (Rhie *et al*. 2021), pooled-sample approaches are still useful for producing draft genomes of very small organisms that may not yield sufficient DNA for high-coverage long-read sequencing (Goldberg *et al*. 2024)

On the one hand, erythrocyte nucleation in plethodontids (Mueller *et al*. 2008) provides for easier extraction and isolation of high molecular weight DNA fragments, facilitating longer read lengths from small amounts of blood. In contrast, many plethodontids are extremely small, as little as ∼25mm SVL (Parra-Olea *et al*. 2016), making it difficult to recover sufficient DNA while retaining an intact specimen. Here, we use HiFi reads on the Pacific Biosciences Revio platform to generate a nearly complete assembly (∼16GB) with high coverage (∼24x) for the Northern Dusky Salamander (Plethodontidae: *Desmognathus fuscus*), from blood and liver collected from two individuals from the same population. We produced 19,640 contigs with an N/L50 of 2,455/1.75MB, the longest of which was 27.9MB, 4,926 were >1MB, and 17,064 were >50KB, representing 99.5% of the genome. A BUSCO search of the genome assembly and kidney/liver transcriptomes revealed ∼93% completeness for the 5,310 tetrapod reference loci.

The relatively small individuals we sampled reduced our ability to produce additional structural data such as chromatin conformation capture, so chromosomal assembly was limited. Scaffolding to the *Ambystoma* genome revealed at least 5.8GB of putatively homologous regions from 3,054 contigs across all 14 chromosomes, but with relatively low contiguity and a high apparent instance of repeat matching across multiple target and query regions. Annotation of repeat elements showed that ∼76% of the genome is repetitive, with ∼35% representing a relatively recent expansion of LTRs (of which ∼66% are Ty3/*gypsy*) and ∼16% representing correspondingly ancient LINE activity. As little as ∼26% of the genome is single copy, in line with previous estimates (Morescalchi and Olmo 1982). Extensive opportunities remain for annotated chromosome-level assemblies of *Desmognathus*, plethodontid relatives, and other salamander families. In the interim, this assembly for *D. fuscus* provides a foundational reference and comprehensive basis for future studies of salamander genomics.

## Materials & Methods

### Sampling & Sequencing

One of us (RAP) collected two *Desmognathus fuscus* (RAP3214–5) from the *‘fuscus* B3’ lineage (Pyron *et al*. 2022; Pyron and Beamer 2023b) on 20 May 2023 in Bell Branch, a small creek on the Thurmond Chatham Game Land in Wilkes County, North Carolina (Fig. 1). The specimens were a relatively small subadult (∼32mm SVL) and adult (∼43mm) captured within ∼3–4m of each other under rocks or logs on the stream bank. Voucher specimens were deposited at the American Museum of Natural History (AMNH A-194068–9). Collecting and research were permitted under NCWRC 23-SC00374 and GW IACUC A2022-004.

**Figure 1.**
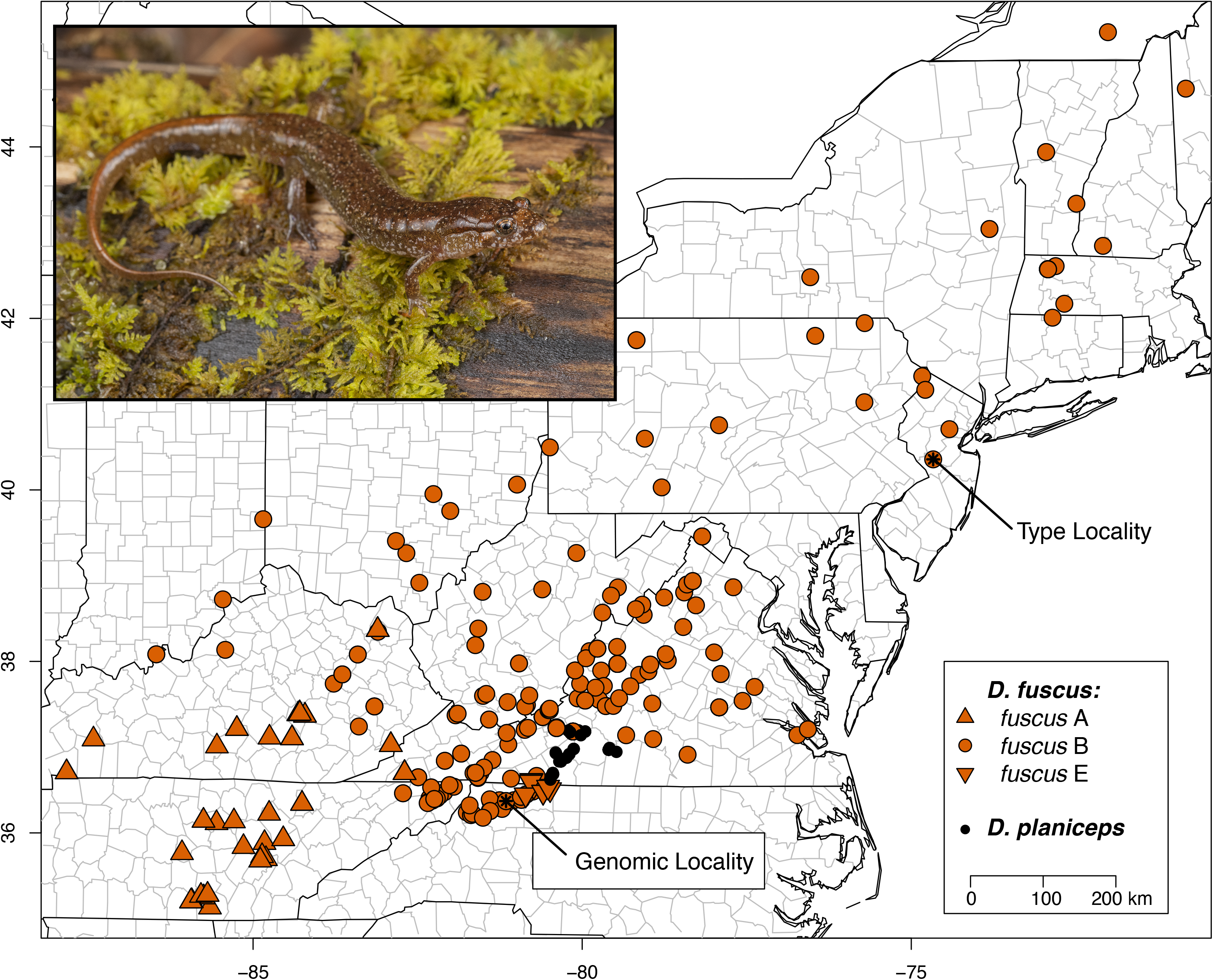
Distribution map of *Desmognathus fuscus* and its sister species *D. planiceps* based on recent taxonomic revisions (Pyron and Beamer 2023b), indicating the type locality (Green 1818; Pyron and Beamer 2020; Dubois *et al*. 2022), and the site in North Carolina where AMNH A-194068–9 were sampled (genomic locality). This species contains phylogeographic sublineages (A, B, and E – indicated by marker shape) which may prove to be distinct species in the future; the B lineage sampled here is nominotypical. Inset photo shows another typical individual in life from the same population (photo courtesy of Max A. Seldes, GW).

Following euthanasia, we extracted blood and liver from both specimens and kidney and associated ducts from AMNH A-194069 only. These were divided among preservation methods, with portions of each retained fresh, flash-frozen at -80C, and preserved in RNALater. We used an NEB Monarch HMW DNA Extraction Kit for Cells & Blood (T3050S) with minimal shearing or fragmentation to purify ultra-high molecular weight genomic DNA from fresh blood. Additionally, we sequenced total RNA transcriptomes from liver (AMNH A-194068) and kidney (AMNH A-194069) at Psomagen Inc. (Rockville, MD) using the TruSeq Stranded Total RNA with Ribo-Zero Gold Human/Mouse/Rat kit. For each sample, we sequenced ∼40M 150bp paired-end reads on a NovaSeq6000 S4 flow cell. We used fastQC (Brown *et al*. 2017) as an initial quality-control screen on the raw reads.

We initially extracted DNA from both specimens, but the larger individual (AMNH A-194069) yielded lower quality. For AMNH A-194068, initial QC at the University of Maryland Institute for Genome Sciences (IGS; http://www.marylandgenomics.org/) using an Agilent Femto Pulse system showed 6,319ng of DNA at 71ng/ul in 89ul of buffered extract (average size 80kb, 85% >= 30kb), which was sufficient for 2 PacBio Revio SMRT Cells. We subsequently provided flash-frozen liver tissue from AMNH A-194069 to the sequencing center for additional UHMW DNA extraction prior to sequencing. This yielded 22,947ng of DNA at 81ng/ul in 284ul, with an average size of 103kb and 79% > 30kb. For each specimen, the IGS performed one PacBio HiFi Library Preparation with standard size selection of 15–18kb, which supplied input libraries for 2 PacBio Revio SMRT Cell 25M Sequencing runs (HiFi/CCS mode, 24 hour movie).

For AMNH A-194068, the two cells yielded 88,311,801,332/85,324,229,953 bases from 6,029,769/5,956,003 CCS reads respectively. This represents a total of ∼174GB of data representing ∼12x total coverage of the expected ∼15GB *Desmognathus* genome (Liedtke *et al*. 2018). The read N50 was 135,250/117,250bp for polymerase reads and 14,754/14,413bp for CCS reads, with a range of 107–57,921bp for the latter. For AMNH A-194069, the two cells yielded 99,120,795,754/92,637,552,538 bases from 5,467,7685,015,293 CCS reads respectively, for a total of ∼192GB of data representing ∼13x coverage. These are slightly better results than for the smaller specimen, possibly representing a difference in the extraction quality or tissue type of blood versus liver. The read N50 was 124,250/113,250bp for polymerase reads and 19,606/16,869bp for CCS reads, with a range of 49–60,229bp for the latter

### Assembly and Scaffolding

We assembled the HiFi reads using hifiasm 0.19.8-r603 (Cheng et al. 2021). Given the low coverage of the individual specimens, we altered the default settings by setting the expected haploid genome size to 14GB (‘--hg-size 14’) and the minimum histogram count for estimating *k-*mer coverage to 2 (‘--min-hist-count 2’). Under these settings, we estimated a diploid assembly for each specimen separately. Individual coverage was not high enough (∼12x) for well-resolved partially phased haplotypes (initial results were highly fragmented), so we instead present the primary (haplotype-merged) assembly for each individual.

Increasing coverage has been shown to result in dramatically higher-quality assemblies up to a threshold of ∼30x, at least in some species with smaller (∼1.6GB) genomes (Xie *et al*. 2023). Accordingly, we then merged the reads from the two individual specimens (∼366GB total) for a high-coverage (∼24x) merged assembly, treating the genome as polyploid (‘--n-hap 4’) to account for the possibly divergent or heterozygous haplotypes across the two individuals. For assemblies that appeared to contain excess contigs from incomplete resolution of highly repetitive regions, we used *purge_dups* to eliminate overlaps (Guan *et al*. 2020). For the transcriptome assembly, we analyzed the reads from both individuals (kidney for AMNH A-194068 and liver from AMNH A-194069) individually in Trinity 2.15.1 (Grabherr *et al*. 2011) to assemble transcript sequences and combined them for an overall transcriptome assembly. After assembly, we filtered the transcripts for quality and residual adapter sequence using bbduk in the bbmap package (https://sourceforge.net/projects/bbmap/).

We assessed initial gene completeness on the genome and transcriptome assemblies using compleasm 0.2.5 (Huang and Li 2023) using the tetrapoda_odb10 reference database (Manni *et al*. 2021b, 2021a). We used the EarlGrey pipeline to identify and annotate transposable elements in the merged primary assembly (Baril *et al*. 2024). We also scaffolded the merged primary assembly to the *Ambystoma mexicanum* reference genome AmbMex60DD (GCA_002915635.3) using RagTag v2.1.0 (Alonge *et al*. 2022). Finally, we extracted, assembled, and annotated the mitochondrial genomes from the individual assemblies using the ‘MitoHiFi’ pipeline (Uliano-Silva *et al*. 2023) with the ‘MitoFinder’ annotation tool (Allio *et al*. 2020) and ‘ARWEN’ for tRNA detection (Laslett and Canbäck 2008). Analyses were performed on the SI High Performance Computing Cluster ‘Hydra’ (https://doi.org/10.25572/SIHPC) and the GW HPCC ‘Pegasus’ (MacLachlan *et al*. 2020).

## Results and Discussion

For AMNH A-194068, the two partially phased haplotypes were 15.5/13.7GB, and the primary assembly was 19.0GB, likely reflecting incomplete resolution of highly repetitive regions given the lower coverage. A single round of *purge_dups* yielded a purged primary assembly of 15.9GB in 51,453 contigs with an N/L50 of 10,433/466Kb and an N/L90 of 1,518/1.02MB. The longest contig is 6.87MB, with 1,619 contigs >1Mb, and 45,297 >50Kb, representing 98.7% of the genome. Similarly, for AMNH A-194069, the two partially phased haplotypes were 16.2/14.2GB, and the primary assembly was 19.2GB. A single round of *purge_dups* yielded a purged primary assembly of 15.8GB in 29,360 contigs with an N/L50 of 5,466/864Kb and an N/L90 of 774/1.97MB. The longest contig is 12.5MB, with 4,215 contigs >1Mb, and 27,385 >50Kb, representing 99.6% of the genome. We named these assemblies *Dfus1*.*0* and *Dfus2*.*0*.

The initial merged attempt yielded a primary assembly of 24.0GB in 65,452 contigs with an N/L50 of 5,306/1.0MB and an N/L90 of 279/5.1MB, suggesting both a dramatic increase in contiguity with the greater pooled sequencing depth (∼24x over ∼12x), and a substantial amount of duplicated haplotigs (∼9GB) in the ‘polyploid’ assembly. Two rounds of *purge_dups* yielded a purged primary assembly of 16.1GB in 19,640 contigs with an N/L50 of 2,455/1.75MB and an N/L90 of 282/5.1MB. The longest contig is 27.9MB, with 4,926 contigs >1Mb, and 17,064 >50Kb, representing 99.5% of the genome. A 16.1GB assembly may still be larger than the true size, representing incomplete merging of repetitive regions, but is similar to previous C-value estimates of ∼13–17pg in the genus based on DNA-Feulgen cytophotometry (Hally et al. 1986). Under Vertebrate Genome Project naming rules (Rhie et al. 2021), we labeled this *aDesFus1-2*.*1* and treat it as our primary assembly for *Desmognathus fuscus* (Fig. 2a).

**Figure 2.**
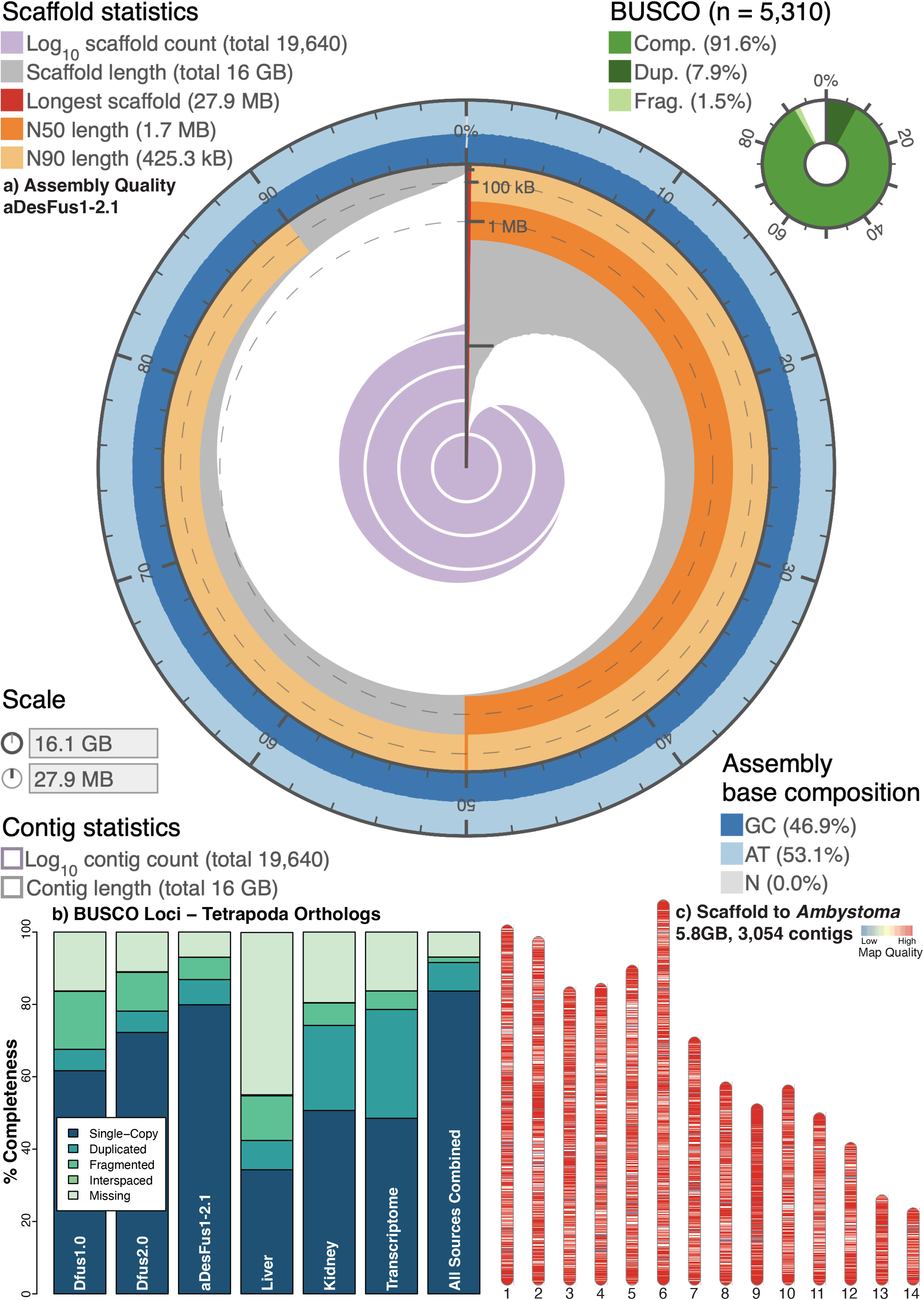
(a) A snail plot (Challis *et al*. 2020) showing the assembly statistics for aDesFus1-2.1 (JBBULT000000000) from AMNH A-194068–9. (b) BUSCO results from compleasm for the various genome and transcriptome assemblies. (c) Scaffolding results from RagTag for the primary merged assembly mapped to the *Ambystoma mexicanum* genome (Schloissnig *et al*. 2021), representing only 5.8GB (∼39%) from 3,054 of 19,640 contigs (∼16%).

Scaffolding to the *Ambysoma mexicanum* genome AmbMex60DD (GCA_002915635.3) placed 5.8GB from 3,054 contigs in 107 scaffolds. These were based on preliminary mappings which, filtered for length (>250bp) and quality (>40), yielded only 405Mb of matching bases from 13,325 *Desmognathus* contigs at 1,319 positions in the *Amybstoma* genome, representing 521,975 possible alignments (Fig. 2c). The longest of these is a copy of the 45S ribosomal RNA tandem repeats (Sochorová *et al*. 2018), with up to 10.3KB of matching bases spread across up to 49.4Kb of alignment to the target region on the long arm of chromosome 6. Two more copies on another contig map to the same target region with similarly high quality.

This low match rate and high variability in repeat matching suggests substantial syntenic divergence across salamanders, given the ∼150Ma split between *Desmognathus* and *Ambystoma* (Shen *et al*. 2016). The other two available assemblies in Salamandridae span the same divergence. Despite this variance, both *Ambystoma* and *Desmognathus* (along with most plethodontids) share a diploid karyotype of 2n=28 (Hally *et al*. 1986; Keinath *et al*. 2015).

Combined with significant recent genome miniaturization in *Desmognathus* (Liedtke *et al*. 2018), we hypothesize that Plethodontidae more broadly and this genus specifically likely harbor unique and highly derived genomic features at the sub-chromosomal level. Consequently, we did not utilize this scaffolding arrangement further in the assembly process.

For the transcriptomic data, fastQC confirmed high quality of the raw Illumina reads, which were filtered and trimmed by trimmomatic (Bolger *et al*. 2014) in the Trinity pipeline. Trinity assembled 246,287 “genes” and 318,347 transcripts from the two tissues, with a contig N50 of 621bp, a median contig length of 297bp, and an average contig length 526bp from 129Mb of assembled data. For the merged genome assembly, compleasm identified 4,941 loci out of 5,310 tetrapod orthologs (93.1%), of which 4,243 were single copy (79.9%), 370 were duplicated (6.97%), and 328 were fragmented (6.18%). The transcriptome contained 4,447 loci (83.8%), of which 2,577 were single-copy (48.5%), 1,597 were duplicated in at least one tissue (30.1%) and 131 duplicated in both (2.5%), and 273 were fragmented (5.1%). Additional filtering for isoforms may reduce the duplication rates in the transcriptome.

Adjusting loci that were fragmented in the genome but single-copy or duplicated in the transcriptome (ignoring fragmented or missing loci in the individual tissues) yielded 4,443 single-copy (83.7%), 420 duplicated (7.9%), and 78 fragmented loci (1.5%) for a total of ∼93% completeness (Fig. 2b). Finally, the annotated mitochondrial genome assemblies have been deposited on Genbank. They match the existing reference (AY728227) for this species (Mueller *et al*. 2004) almost exactly in terms of length, gene order, and base composition. In coding or functional regions, they differ from each other by a 2bp indel in the 16S ribosomal RNA gene, a single transition in cytochrome *b*, and two nearby indels in the putative control region/*D*-loop.

We also performed an additional compleasm run on the ∼5.8GB that can be roughly scaffolded to the *Ambysoma* genome, to assess the extent to which these putatively homologus regions were enriched for the core set of tetrapod orthologs. The query matched 2,506 (47%) of the 5,310 reference loci, suggesting that roughly half of the BUSCOs in the assembly are concentrated in moderately conserved regions. In contrast, almost 50% occur in regions too divergent to easily map across the 150Ma divergence between the two lineages.

Analysis of transposable elements in the primary assembly yielded several interesting patterns (Fig. 3). Approximately 26% of the genome is single copy, while ∼74% is repetitive (Fig. 3c), greater than that of *Ambytoma* at ∼68% and similar to that of *Pleurodeles*, ranking among the highest percentages known in animals (Meyer *et al*. 2021; Schloissnig *et al*. 2021; Brown *et al*. 2022). However, *Desmognathus* is more similar to *Ambystoma* in having ∼35% of the genome representing LTRs and ∼15% representing LINEs, whereas the majority of *Pleurodeles* TEs are DNA-based. In contrast, DNA transposons represent only ∼3% of the *Desmognathus* genome, substantially less than *Ambystoma* and *Pleurodeles*. Unlike those two species for which relatively few TEs were unknown, ∼17% of the *Desmognathus* TEs are unclassified (Fig. 3a). This may be an artifact of our unscaffolded assembly or the identification steps in the EarlGrey pipeline, which we did not optimize for amphibians. Alternatively, there may be additional novel elements or artifacts associated with *Desmognathus*, potentially related to the significant down-shift in genome size that occurred in the ancestral lineage (Liedtke *et al*. 2018).

**Figure 3.**
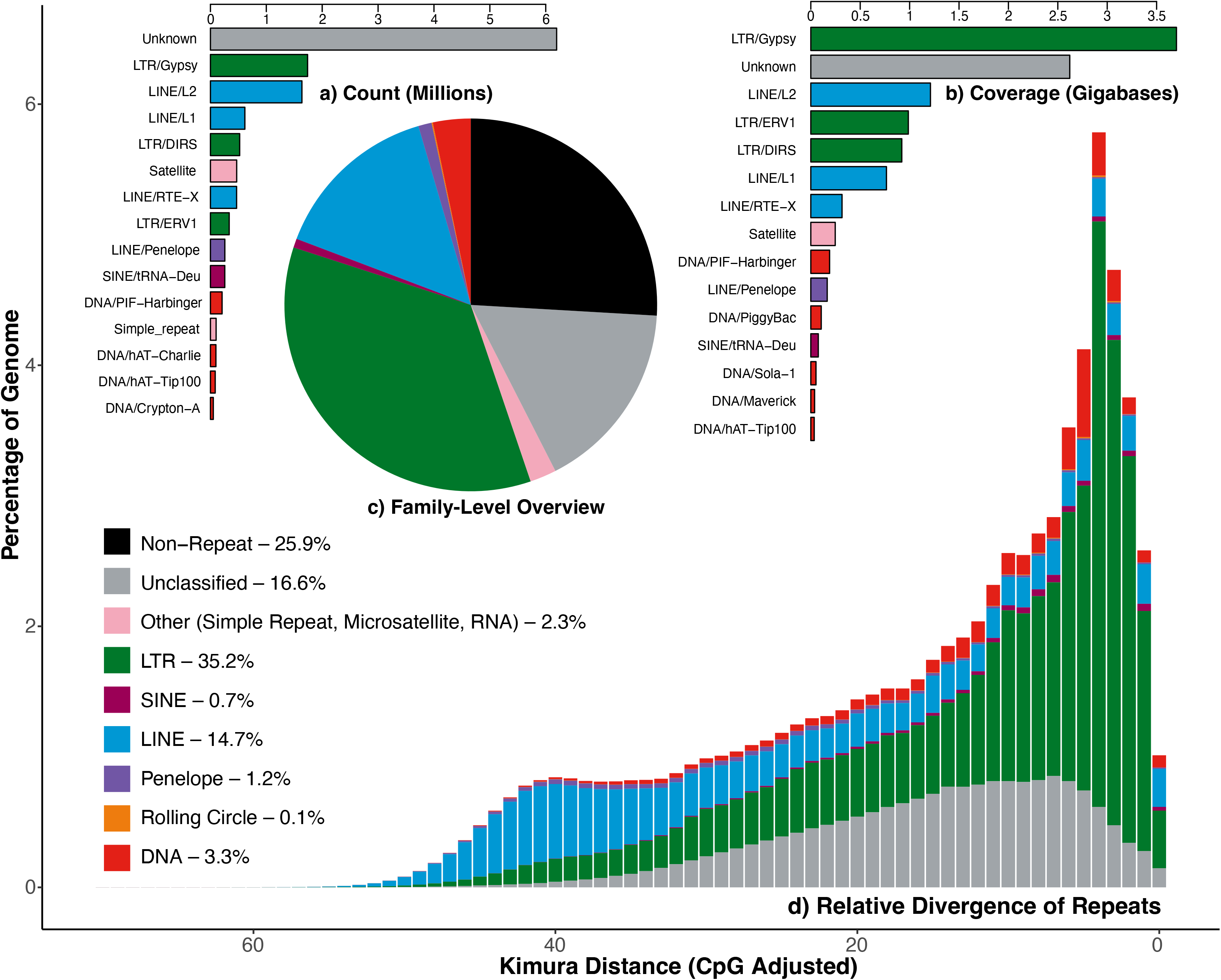
The transposable element landscape for aDesFus1-2.1 identified and annotated using the EarlGrey pipeline (Baril *et al*. 2024), showing TEs by count > 50k (a), coverage > 30MB (b), percentage by family (c), and relative K2P divergence (d). The divergence landscape (showing oldest to most recent, left to right) indicates an early proliferation and slow contraction of LINEs, followed by recent massive LTR expansion and smaller but substantial increases in unclassified elements and DNA transposons, with relatively little very recent or non-divergent activity.

Similar to earlier shotgun-based estimates for plethodontid and cryptobranchid salamanders (Sun *et al*. 2012b; Sun and Mueller 2014) and the more complete recent genomes for ambystomatids and salamandrids (Schloissnig *et al*. 2021; Brown *et al*. 2022), Ty3/*gypsy* LTRs are the dominant TE in *Desmognathus*, comprising ∼3.7GB or 32% of repeats (Fig. 3b).

Others such as LINE/L2, LTR/DIRS, and LTR/ERV1 are similarly common across salamanders. Unlike *Ambystoma* and *Pleurodeles* where PIF/Harbinger elements are ∼7–20% of repeats, they are much less common (<3%) in *Desmognathus* and other plethodontids. A novel finding in our analysis is a relatively high percentage of satDNA (2.1% of repeats) compared to *Ambystoma* and *Pleurodeles* (<0.5%); more similar to the ∼5% in *Homo* (Brown *et al*. 2022).

In our *Desmognathus fuscus* assembly, the relative divergence landscape shows an early expansion and slow contraction of LINEs, followed by a quick recent expansion of both LTRs and DNA transposons (Fig. 3d). Similarly, the large group of unclassified elements expanded at roughly the same time, and very little of the genome consists of more recent or non-divergent repeat activity. The interplay between TEs, slow DNA loss, and genome expansion is well known in salamanders (Sun and Mueller 2014). How these dynamics interacted and contributed to the dramatic *C-*value decrease in *Desmognathus* (Liedtke *et al*. 2018; Sclavi and Herrick 2019) represents a pertinent question for future research.

A previous study sampling four species spanning the range of sizes, ecotypes, and phylogenetic lineages in the genus (*Desmognathus amphileucus, D. monticola, D. perlapsus*, and *D. wrighti*) showed a uniform diploid karyotype of 2*n*=28, suggesting that this structure is conserved across the group (Hally *et al*. 1986). While most authors have reported *C-*values of 14–16pg for *Desmognathus* species (Hally *et al*. 1986; Sessions and Larson 1987; Liedtke *et al*. 2018), some studies have reported values up to 18–22pg for species including *D. fuscus* (Bachmann 1970; Olmo 1974). Methods for determining genome size vary among these studies (e.g., Feulgen densitometry, flow cytometry), potentially explaining some of the variation. As the taxonomy of *Desmognathus* and *D. fuscus* in particular have shifted dramatically in recent years (Pyron and Beamer 2023b), comparisons between previous results are difficult since sampling localities were often not reported for measured specimens. High-coverage assemblies with chromosome-level scaffolding from *D. fuscus* and other species are needed for an accurate assessment of genome-size variation in the group.

We note that one previous study intentionally testing the life-history/genome-size correlation in *Desmognathus* rejected the hypothesis, finding no relationship (Hally *et al*. 1986). In contrast, the largest *C-*values of 18–22pg (Bachmann 1970; Olmo 1974) were estimated for *D. “marmoratus”* and *D. “quadramaculatus*,*”* (see Pyron and Beamer 2022, 2023a for recent taxonomic revision of these complexes) which are both highly aquatic and lay their eggs in water, the ecological state associated with larger genomes in other salamander lineages (Lertzman-Lepofsky *et al*. 2019). Similarly, the smallest *C-*values (<14pg) are associated with the terrestrial direct-developer *D. wrighti* (Hally *et al*. 1986; Sessions and Larson 1987). This and similar results for other terrestrial direct-developing Neotropical plethodontids recently discovered to have miniaturized genomes (Decena-Segarra *et al*. 2020) suggests the potential for complex evolutionary and ecological interactions across Plethodontidae.

## Data Availability Statement

The genome and transcriptome assemblies and raw reads for both HiFi (DNA) and Illumina (RNA) sequencing are deposited on SRA under BioProject PRJNA1038779 and BioSamples SAMN38201181–2 for AMNH A-194068–9 (SAMN40589320 for the merged reads). The mitochondrial genomes are deposited on Genbank under accessions OR794553/PP503028. This Whole Genome Shotgun project has been deposited at DDBJ/ENA/GenBank under the accessions JAXCVG000000000 (AMNH A-194068/Dfus1.0), JBBMEE000000000 (AMNH A-194069/Dfus2.0), and JBBULT000000000 (aDesFus1-2.1). The versions described in this paper are versions JAXCVG010000000, JBBMEE010000000, and JBBULT010000000, respectively. The transcriptome assemblies are GKTJ00000000 (AMNH A-194068 - liver) and GKTI00000000 (AMNH A-194069 - kidney).

## Acknowledgments

The authors thank S. Pirro (NCBI), T. Hains (Chicago), K. O’Connell (Deloitte), M. Tollis (NAU), L. Sadzewicz and L. Tallon (UMD), and R. Rautsaw, S. Robins, B. Shin, I. Dzekunova, N. Bittner, and A. Carvalho (PacBio) for recommendations and assistance with genome assembly and annotation. We also thank D. Beamer (ECU) and the North Carolina Wildlife Resources Commission for permits and logistical support. We gratefully acknowledge the computing resources provided on the High Performance Computing Cluster operated by Research Technology Services at the George Washington University, and the Smithsonian Institution High Performance Computing Cluster.

## Abbreviations

BUSCO: Benchmarking Universal Single-Copy Orthologs

## Authors’ Contributions

EAM and RAP contributed to all parts of the MS equally.

